# A Real-Time Automated Deep Learning Workflow for Non-invasive High-Magnification Imaging of *C. elegans*

**DOI:** 10.64898/2026.06.01.729022

**Authors:** Parsa Safaeian, Tasnuva Binte Mahbub, Rhythem Tahrin, Mohammad Tanha, Mark W. Pellegrino, Salman Sohrabi

## Abstract

*Caenorhabditis elegans* is a premier model organism for aging and neurobiology research, valued for its short lifespan, optical transparency, genetic tractability, and well-mapped nervous system. Non-invasive automated recording of biomarkers is a fundamental goal in modern biology because it preserves natural physiology and eliminates confounds from anesthesia, restraint, or repeated handling in *C. elegans*. Yet high-magnification imaging of freely moving worms remains a persistent challenge: as magnification increases, the narrowing field of view compounds target loss, motion blur, and focal drift, pushing researchers toward immobilization strategies that compromise physiology, suppress natural behavior, and preclude the continuous longitudinal observation essential for aging and neurobiological studies. Here, we present a real-time tracking workflow for imaging individual worms in a microfluidic platform under controlled culture conditions. The system integrates deep learning head detection, image-based autofocus, and rapid motorized-stage feedback to support stable imaging across multiple magnifications, including neuronal-scale imaging. Hundreds of individually housed worms in separate incubation chambers enable repeated daily imaging of the same animals throughout their lifespan. Built entirely on a commercially available inverted microscope without additional custom hardware, the platform features a modular, user-configurable interface adaptable to diverse microscope setups, specimens, and experimental goals. Fluorescence images from freely moving worms were visually comparable to those from immobilized animals, supporting longitudinal phenotyping in aging and neurobiology studies.

## Introduction

*C. elegans* is a widely used model organism for studying development, metabolism, neurobiology, reproduction, and aging [1]. This model is especially useful for aging research because of its short lifespan, observable age-associated phenotypes, genetic tractability, and optical transparency [2]. Its high reproductive output, small size, simple culture requirements, and compatibility with high-throughput assays, together with strong evolutionary conservation, also make it well suited for large-scale studies [4,5].

Despite these benefits, C. elegans assays often remain labor-intensive and time-consuming, often requiring trained researchers to perform repeated manual picking, continuous monitoring, and scoring over more than two weeks [3]. This heavy reliance on manual handling not only limits throughput but also increases variability and the risk of subjective error, thereby reducing consistency and reproducibility [4]. These challenges become especially pronounced when daily microscopy and repeated imaging are required over extended periods.

To address this problem, automated imaging platforms have been developed using standard culture formats such as Petri dishes and multiwell plates. Several platforms, including CARIBN II, Lifespan Machine, WorMotel, and more recent multimodal imaging systems, have improved throughput and enabled automated observation of worm behavior, survival, and physiology [5–7]. While these platforms have substantially expanded the capacity for longitudinal phenotyping in C. elegans, plate-based culture imposes fundamental limitations. Distinct experimental conditions must be maintained on separate plates, increasing labor, and each plate typically houses tens of worms, making it difficult to track individual animals over time. Because responses to environmental change can vary across tissues and among individuals, these constraints collectively create a compelling need for single-animal longitudinal phenotyping under well-controlled conditions.

Microfluidic platforms address these needs by enabling precise environmental control, supporting stable long-term culture, and confining individual worms within separate chambers [6–8]. When integrated with robotics and software-driven microscopy, these platforms enable real-time monitoring [8,9]. WormSpa maintained individual worms in separate chambers under constant low flow and enabled continuous monitoring of single-animal responses, such as immune-response gene expression and egg laying [10]. A key limitation was persistent head movement, which reduced suitability for single-cell imaging, especially in head cells. HeALTH increased throughput by integrating automated behavioral imaging with continuous flow control and Peltier-based temperature regulation, allowing monitoring of more than 1,400 isolated worms under tightly controlled spatiotemporal conditions throughout adulthood. This revealed behavioral and lifespan responses to diet and temperature, though the use of low-magnification imaging limited fine-resolution imaging [11]. Lifespan-on-a-chip cultured individual worms from the L4 stage until death and used reversible clamps for brief immobilization, enabling repeated measurement of body size and locomotion across life [12]. Its throughput was limited to about 16 worms per experiment, and the intermittent restraint did not fully reflect natural behavior.

A major challenge that remains in this field is high-magnification imaging of freely moving worms. This limitation becomes especially pronounced when small anatomical structures, such as individual neurons, must be kept within a highly restricted field of view during continuous movement. At magnifications above 20×, the field of view becomes too small to capture the entire worm, necessitating rapid image acquisition, real-time target detection, and fast stage correction to keep the region of interest in view during movement. In contrast, although objectives below 20× reduce target loss by expanding the observable area, their spatial resolution is often insufficient for reliable visualization of individual neuronal somata, particularly within the densely packed head ganglia around the nerve ring [13]. Thus, high-magnification imaging of freely moving animals remains susceptible to motion blur, repeated target loss, and frequent stage corrections. To obtain stable high-magnification images, many studies have relied on worm immobilization. Chemical immobilization with agents such as levamisole or sodium azide is widely used, acting by pharmacologically paralyzing the animals through cholinergic activation or inhibition of mitochondrial metabolism, but these treatments prevent feeding and disturb physiological function [14]. Cooling-based immobilization suppresses movement by lowering temperature, but it can induce stress, increase rupture risk during rapid temperature shifts, and provide only a short imaging window after return to room temperature [15]. A light-controlled method based on photothermal Pluronic F127 immobilization enables reversible immobilization of C. elegans by using illumination-induced heating to trigger a thermoreversible sol-gel transition under near-physiological conditions. It is not instantaneous and depends on careful thermal control [16]. Physical stabilization with polystyrene microbeads can support extended live imaging, but it does not fully eliminate residual head and tail movement, and recovery can be complicated by drying [17]. Microfluidic confinement immobilizes C. elegans by trapping worms in narrow channels or with compression or suction to limit movement during imaging; however, it relies on physical restraint, specialized fabrication, and often pressure-control hardware, which can affect physiology and complicate long-term or recovery-based experiments [18]. Hydrogel encapsulation can be gentler than direct mechanical restraint, but temperature-responsive gels may still impose cold-related stress, whereas photo-crosslinked hydrogels can introduce UV-induced cellular damage [19,20]. Critically, none of these immobilization strategies is compatible with sustained incubation over the animal’s full lifespan, as repeated immobilization events accumulate physiological stress, restrict natural behavior, and ultimately compromise the biological validity of longitudinal measurements.

To address these limitations, we developed a tracking workflow for non-invasive longitudinal imaging of freely moving C. elegans in a reusable microfluidic chip under controlled culture conditions. The platform supports multi-scale imaging at 10×, 20×, and 40× magnifications, enabling applications ranging from whole-animal observation to high-resolution imaging of fine anatomical features, including neurons. The workflow integrates deep learning-based detection and segmentation with rapid stage feedback and image-based autofocus to maintain the worm head within the field of view during continuous movement. Operating at 22 fps with a feedback cycle of approximately 45 ms, the system supports rapid updates during detection, recentering, and tracking. Individual housing within the microfluidic chip preserves worm identity while maintaining a regulated environment over time and allowing unrestricted motion during longitudinal observation. We validated the system at 40× magnification, where imaging constraints are most severe, and found that workflow performance was sensitive to the detection confidence threshold, with intermediate values providing the most effective balance between tracking initiation and continuation. Image acquisition is triggered only after the animal satisfies various criteria, thereby reducing motion blur and improving fluorescence image quality. The platform acquires multiple high-resolution images per worm per day. By integrating single-animal microfluidic culture, AI-based real-time recentering, autofocus, and behavior-triggered image acquisition within a single workflow, this platform enables non-invasive high-magnification longitudinal imaging of freely moving worms under unrestricted motion, while yielding image quality comparable to that obtained from immobilized animals.

## Results

An adult C. elegans hermaphrodite is approximately 1 mm in length and 70–80 µm in width [21]. At 40× magnification, the 400 × 400 µm field of view is narrow relative to the speed of worm locomotion, which may approach 800 µm/s. In practical terms, even a head initially centered in the frame may leave the imaging area within about 250 ms, leaving only a narrow window for reliable tracking and recentering by the motorized stage. This constraint becomes even more pronounced when imaging neuronal structures, because the dimensions of individual somata are small relative to the displacement that can occur during even a short exposure of 25 ms, resulting in substantial motion blur. During a single exposure, a neuron may move by several cell diameters, spreading its signal over a larger pixel area and reducing the signal-to-noise ratio at individual pixels. To address this problem, we developed a high-speed tracking system that continuously identifies the worm head and adjusts the motorized stage in real time to keep the target within the field of view during imaging. Operating at approximately 22 frames per second, the system completes image processing and rapid positional updates in about 45 ms per cycle. This update rate is sufficiently fast to support continuous tracking during high-magnification imaging. The chip also contains over 300 individual incubation chambers. At full capacity, the platform typically acquires about 2 to 5 images per worm per day. When imaging is focused on a smaller cohort, such as about 100 worms, the sampling rate increases to approximately 7 images per worm per day. A major strength of this system is its ability to support repeated daily imaging throughout the animal’s lifespan without adverse biological effects. By eliminating the need for immobilization and enabling rapid tracking, the system supports longitudinal monitoring while preserving natural behavior.

### Tracking System Overview

High-magnification tracking of C. elegans is performed using an integrated microscopy system (Figure 1A) comprising an inverted microscope, a high-speed camera, and a precision motorized XY stage. Microscope operation, imaging automation, and stage motion are managed through a LabVIEW supervisory interface that executes Python modules for tracking and image processing. This framework enables automated control of microscope components, including the motorized z-drive, motorized objective turret, contrast modes, and illumination settings, thereby supporting automated 2D worm tracking in the XY plane with axial focus control at 10×, 20×, and 40× magnification. Worms are housed in a microfluidic chip containing 384 individual chambers, each designed to confine a single animal. Image acquisition is performed using an sCMOS camera operated at up to 664 fps with 8-bit image depth. The motorized XY stage enables precise, high-speed motion and provides submicron positioning control, with travel speeds of up to 500 mm/s, 5 nm resolution, and positioning accuracy of less than ±1.0 µm.

**Figure 1.**
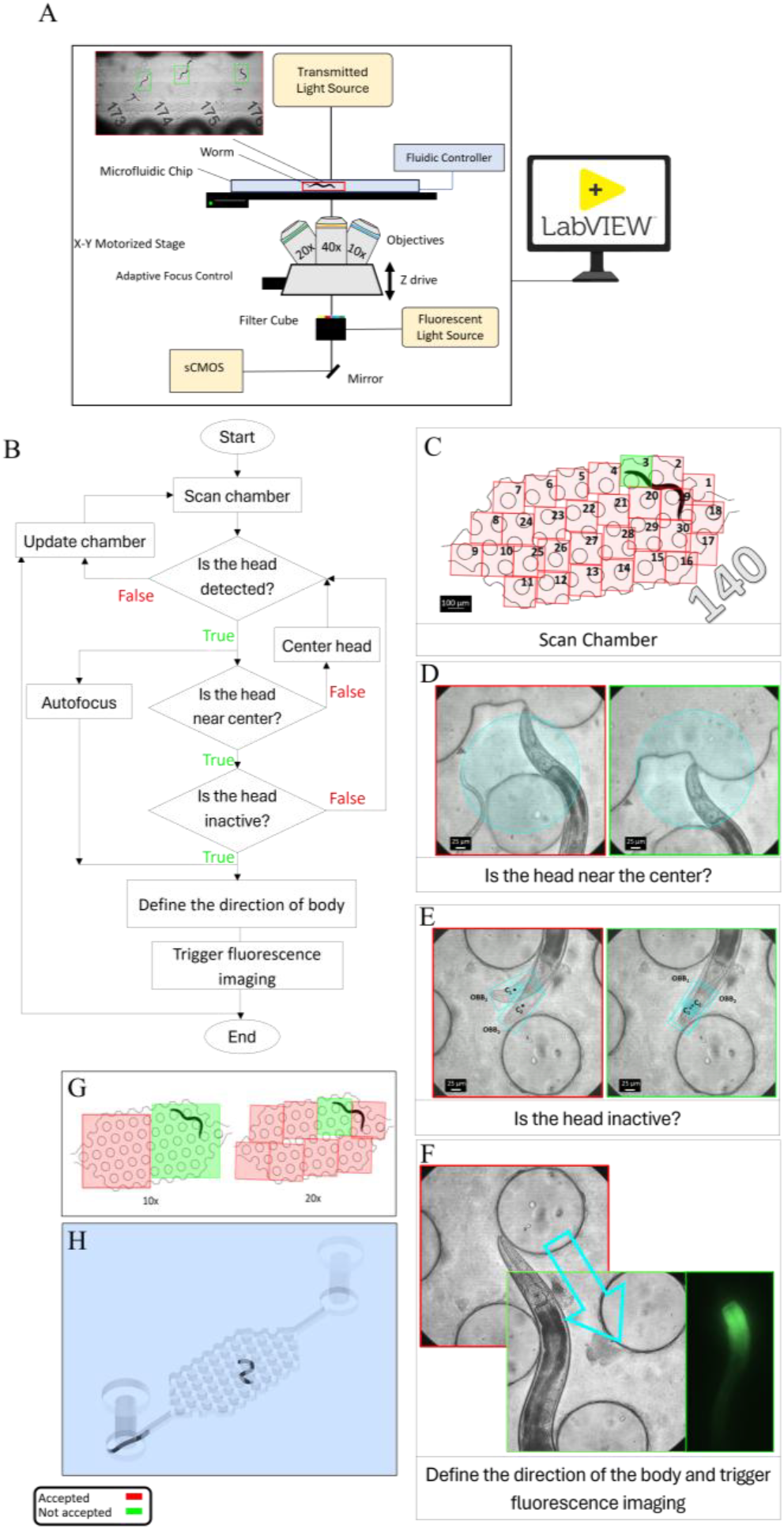
**(A)** Schematic of the imaging platform, including an sCMOS camera, fluorescence and transmitted-light illumination, filter cube, Adaptive Focus Control (AFC), objectives, motorized stage, and z-drive, all operated through a LabVIEW interface. **(B)** Tracking workflow. Each chamber was first scanned to localize the worm, after which the detected head was used to initiate tracking. The stage then recentered the worm during movement while autofocus established the optimal focal plane. Tracking continued until the worm satisfied the inactivity criterion, which triggered image acquisition. **(C)** Schematic of the tiled scan sequence used to locate a worm within chamber 140 at 40× magnification. Scanning begins at tile 1 and proceeds sequentially through tile 30. In this example, the worm head was not detected in tiles 1 and 2 (red), whereas tile 3 (green) was the first tile in which the head was detected, marking the start of tracking. **(D)** Center Head. The red image shows the worm out of the center, whereas the green image shows the worm head positioned within the circular target region. **(E)** Inactivity-state classification for fluorescence triggering. After autofocus was completed, the first frame was designated as the reference frame, and the detected head in this frame was defined as OBB1 with centroid C1. In each subsequent frame, the newly detected head was defined as OBB2 with centroid C2. If C2 remained within a 50 µm-diameter circular region centered at C1 for four consecutive frames, corresponding to 200 ms, the worm was classified as inactive, and fluorescence imaging was triggered. If this condition was not met, OBB1 and C1 were updated, and the evaluation was repeated. The red image indicates an active state, in which C2 falls outside the predefined circular region, whereas the green image indicates an inactive state, in which C2 remains within the region and satisfies the triggering criterion. **(F)** Movement toward the midbody. The arrow indicates the direction of movement, and the corresponding brightfield and fluorescence images are displayed. **(G)** Schematic of the tiled scan pattern used to find the worm within a chamber at 10×and 20×. **(H)** Schematic of worm position showing one worm inside the chamber and another worm within the connecting channels.

### Tracking Workflow

The dimensions of each chamber in our microfluidic chip are 3800 × 1700 µm. In contrast, the field of view at 40× magnification is only 400 × 400 µm, preventing the entire chamber from being captured in a single frame. The first step is to detect worms within each chamber (Figure 1B). A rapid tiled scan is acquired and processed using a YOLO-based head-detection model (Figure 1C). The number 140 indicates the chamber ID. If no worm is detected, the system advances to the next chamber (e.g., chamber 141) and continues scanning. Once a worm is detected, the system locks onto the head, and tracking begins. During tracking, the motorized stage repositions the field of view to keep the detected worm’s head centered (supplementary Movie S1). In general, recentering is triggered when the worm head approaches the boundary of the field of view, using a circular target region with a diameter of approximately 300 µm, corresponding to about 70% of the frame width (Figure 1D). This conditional recentering strategy minimizes unnecessary stage movements and reduces motion blur during imaging, as continuous unconditional recentering requires frequent high-speed corrections that can destabilize tracking. Concurrently, autofocus is executed to determine the optimal z position for the worm, during which the x, y, and z coordinates are updated. After focus is established, the z position remains constant until the worm enters an inactive state suitable for image acquisition. Inactivity is defined as the center of the detected head oriented bounding box (OBB) remaining within a 50 µm-diameter circular region for 200 ms (Figure 1E), indicating only small local oscillations. This threshold is comparable to the approximate width of the head-body region of a young adult C. elegans. Once inactivity is confirmed, fluorescence imaging is triggered. For head imaging, no additional stage movement is required. For mid-body imaging, however, the stage is repositioned to align the worm’s orientation with the target body region in the field of view (Figure 1F).

Importantly, the workflow is compatible with multiple objective magnifications, with the main difference being the scanning strategy. At lower magnifications, the larger field of view reduces the number of tiled frames needed to cover an incubation chamber, whereas higher magnifications require denser tiling. Because the field of view scales approximately with 1/M^2^, the number of tiled frames increases with magnification. In our system, complete chamber coverage required 2 frames at 10×, 8 frames at 20×, and 30 frames at 40× (Figure 1G).

#### Focus Optimization

Accurate focus control is essential for high-resolution microscopy, particularly in automated imaging systems where samples may move during acquisition [19]. In microfluidic imaging platforms, small variations in device geometry, chamber height, and specimen position can shift the optimal focal plane across the field of view, making dynamic focus adjustment necessary to maintain consistent image sharpness. Hardware-based methods such as Adaptive Focus Control (AFC) provide a stable initial focal reference from reflective optical interfaces, whereas image-based autofocus refines focus directly from specimen image quality. In this study, AFC, an infrared-based hardware autofocus system, was used during chamber-to-chamber movement to establish an initial focal reference [19]. AFC maintains the distance between the objective and the bottom surface of the glass slide, but this alone is not sufficient for high-magnification imaging because it cannot compensate for variations in slide thickness or specimen movement along the z-axis. As a result, the AFC reference plane did not always coincide with the worm’s true plane of best focus during tracking, so focus was further refined using image-based sharpness evaluation.

Our first step toward addressing this issue was to develop an image-processing strategy for identifying the focal plane. To compare sharpness across different axial positions, we used images of an immobilized worm acquired at multiple focal planes. Immobilization ensured that image features remained unchanged between acquisitions, allowing direct comparison of sharpness measurements. The in-focus position was defined as z = 0, and axial image stacks were collected from −100 to +100 µm relative to the in-focus plane without any x-y movement of the stage. In the initial analysis, image sharpness was computed over the entire frame using three gradient-based metrics: Tenengrad, Sobel variance, and Laplacian variance. However, the resulting sharpness profiles exhibited multiple local maxima rather than a single distinct peak, making it difficult to determine the true focal plane (Figure 2A–C). Sharpness values also varied inconsistently across axial positions and did not reliably reflect the worm’s actual focus, such that higher sharpness values did not always correspond to a sharper image of the animal. This inconsistency was primarily caused by background structures within the field of view, including microfluidic channel features, debris, eggs, and larvae.

**Figure 2.**
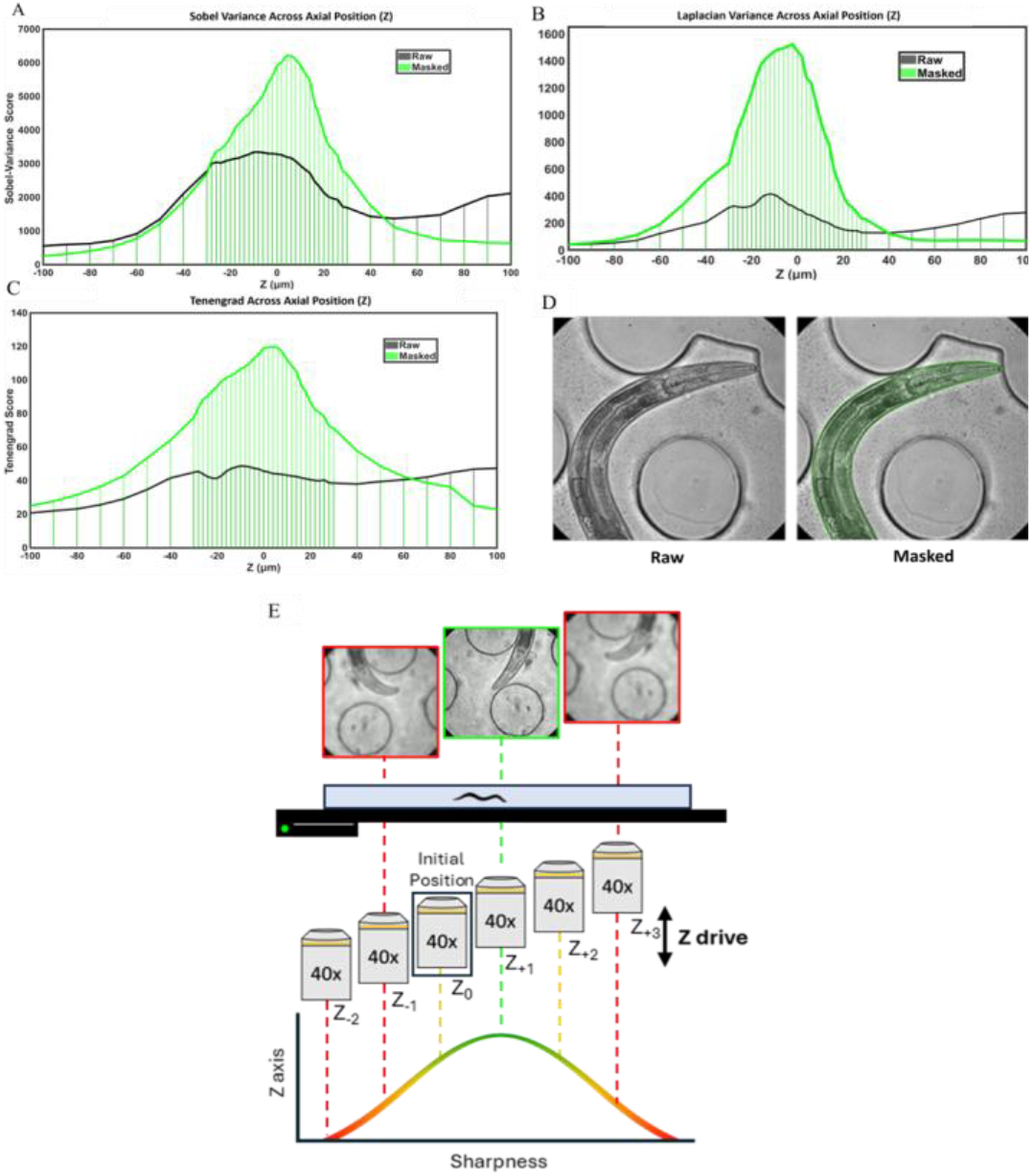
**(A–C)** Effect of worm masking on sharpness–z profiles calculated using Laplacian variance, Sobel variance, and Tenengrad. Sharpness values from the raw image are compared with those from the masked worm region. Masking improves peak definition and best-focus localization by reducing background contributions. Across all three metrics, the masked curves (green) peak near the user-defined best-focus plane (z = 0), with Sobel variance and Tenengrad peaking at z = +4 µm and Laplacian variance peaking at z = −2 µm. **(D)** Representative raw and masked images. **(E)** Autofocus Example. Starting from the AFC-defined focal position, the motorized z-drive moves above and below the initial z position using small steps, which can be as low as 2 µm. Each image is segmented, and sharpness is measured only within the worm region. In the example shown, +1 step gives the highest sharpness and is selected as the candidate focal plane. This is supported by the −1 and +3 step images, which are blurrier and have lower sharpness values. Sharpness increases as the focal plane gets closer to the best-focus position and decreases as it moves away. Only a few sampled positions are shown here as an example. In practice, the autofocus routine can scan more than 40 steps, covering a total z-range of more than 80 µm when a 2 µm step size is used.

**Figure 3.**
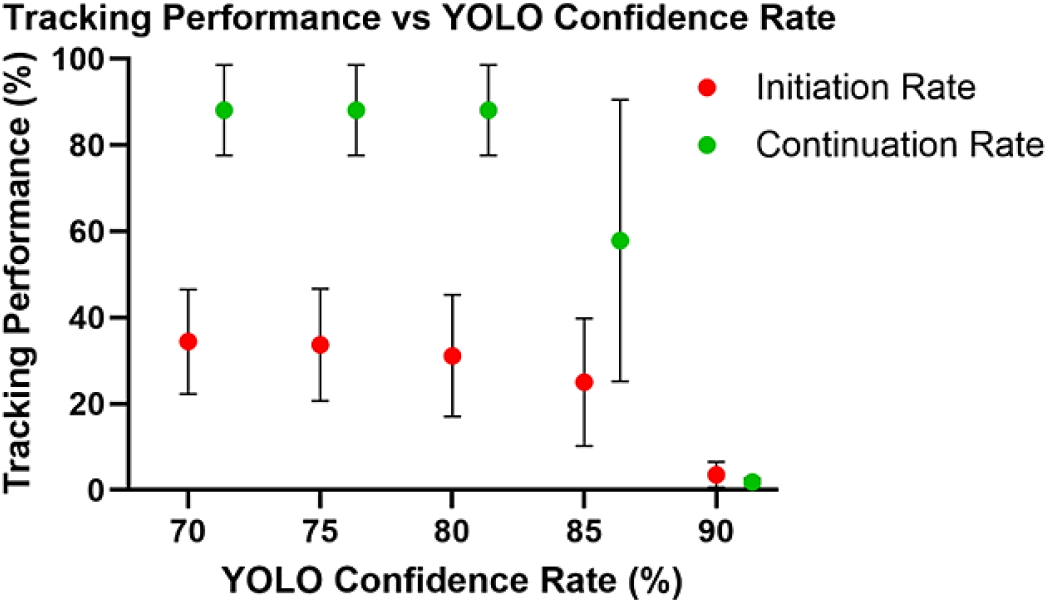
Effect of YOLO confidence threshold on tracking performance (tracking initiation and tracking continuation). At 90%, tracking performance was nearly abolished, with successful outcomes observed in only about 1% of runs. Lowering the threshold markedly improved both metrics, with initiation and continuation reaching 31.1 ± 14.1% and 88.1 ± 10.5%, respectively, at 80%. At 70%, continuation remained unchanged at 88.1 ± 10.5%, whereas initiation increased only modestly to 34.4 ± 12.1%, indicating that thresholds in the range of 80–70% provided the most favorable balance between sensitivity and tracking stability.

To overcome this limitation, the analysis was limited to the worm body region (Figure 2). The segmentation model was used to generate a worm mask, and sharpness was calculated only within the masked region (Figure 2D). By excluding background features from the calculation, interference in the sharpness measurements was eliminated. After masking, all three metrics exhibited a single well-defined peak, enabling reliable identification of the highest-sharpness frame as the focal plane (Figure 2 A–C). In this masked analysis, sharpness was maximal near focus and declined with increasing distance from the focal plane, producing a clear unimodal profile. All three sharpness metrics performed consistently, and the mean axial position of their peak sharpness values was 2 µm, close to the predefined in-focus position at z = 0, supporting the validity of the evaluation method.

After establishing a workflow for identifying the optimal focal plane, the next challenge was to integrate autofocus during active tracking of freely moving animals. To evaluate sharpness without interrupting real-time tracking, sharpness calculations were performed in a separate Python thread, allowing autofocus to run in parallel with the main tracking loop. Starting from the AFC-derived initial z-position (*z*_0_), we used the microscope z-drive to move the focal plane two steps upward (*z*_+1_, *z*_+2_)and two steps downward (*z*_−1_, *z*_−2_). At each z position, an image was acquired, the animal was segmented, and sharpness was calculated only within the masked worm region to reduce background interference from chamber edges, debris, or illumination artifacts (Figure 2E). The algorithm then compared the mask-based sharpness values across the tested planes. If *z*_0_was not the sharpest plane, the system moved in the direction where sharpness increased. For example, if sharpness increased at *z*_+1_, the z-drive moved one additional step upward to *z*_+3_ to verify whether *z*_+1_represented a local maximum. If sharpness decreased at *z*_+3_, *z*_+1_was selected as the optimal focal plane.

### Tracking Performance Optimization

Tracking performance was evaluated using two criteria: tracking initiation and tracking continuation. Tracking initiation success rate (TI) was defined as the percentage of scan events in which the tracking workflow was successfully initiated. An initiation event was considered successful only when a worm was detected during scanning, and tracking could be started. Initiation failure occurred when no worm was detected, for example, because the worm was outside the chamber during scanning (Figure 1H). Scan events with no valid detection were counted as initiation failures.

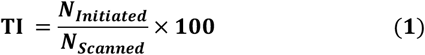

where *N*_Initiated_ is the number of chambers in which tracking was successfully initiated, and *N*_*scanned*_ is the total number of scanned chambers.

Tracking continuation success rate (TC) was defined as the percentage of initiated tracking events in which the worm remained continuously tracked without being lost. This metric quantified how reliably the workflow maintained lock on a worm after tracking had been initiated. Continuation failure occurred when the worm was lost during tracking, for example, due to rapid movement, occlusion, or consecutive head-detection failures.

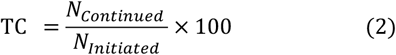

Overall tracking performance was defined as the product of tracking initiation success rate (1) and tracking continuation success rate (2):

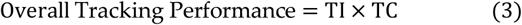

This metric provides a combined measure of the workflow’s ability to initiate tracking and sustain it through image acquisition. Image quality was assessed separately as the fraction of acquired frames that met predefined measurement-specific criteria. A frame was counted as acceptable only if it was sufficiently sharp and the intended anatomical region was fully contained within the field of view.

The YOLO confidence threshold strongly influenced both tracking initiation and tracking continuation. At the most stringent threshold of 90%, both metrics were nearly abolished, with successful outcomes observed in only about 1% of runs. Lowering the threshold to 85% markedly improved performance, and a further reduction to 80% increased initiation to 31.1% and continuation to 88.1%. At 70%, continuation remained unchanged at 88.1%, whereas initiation increased only modestly to 34.4%. Together, these findings indicate that excessively high confidence thresholds substantially reduce detection sensitivity and impair both workflow initiation and sustained tracking, whereas thresholds in the range of 80–70% provide the most favorable balance between sensitivity and tracking stability.

To calculate overall tracking performance, the average initiation and continuation rates were multiplied at each threshold. Using this definition, the highest tracking performance was obtained at 70%, where the mean initiation rate of 34.4% and the mean continuation rate of 88.1% yielded an overall tracking performance of 30.3%. However, if initiation is considered as 100%, such as in an idealized case where the worm is always successfully detected at the start, the maximal effective tracking performance corresponds to the continuation plateau of 88.1%.

To evaluate imaging performance under natural behavioral conditions, brightfield and fluorescence images of the AIY reporter strain OH99 were acquired from freely moving C. elegans shown in Figure 4A and compared with matched images from immobilized animals (Figure 4B). AIY is a pair of head interneurons that integrates sensory information and helps regulate behaviors such as thermotaxis, olfactory responses, and locomotion in C. elegans [22,23]. Despite the absence of chemical or physical restraint, freely moving worms produced image quality comparable to that of immobilized preparations, with clear visualization of overall morphology in brightfield and distinct AIY fluorescence (Figure 4A). The platform also supported imaging at 10×, 20×, and 40× magnifications, with GR1719 shown as an example of lower-magnification imaging. At 10×, the system was well-suited for rapid whole-animal monitoring, including tracking, posture, body size, survival, and global fluorescence measurements. At 20×, it enabled organ- and cell-level analysis while maintaining a sufficiently broad field of view for longitudinal sampling, including intestinal reporter quantification and measurement of variation in nuclear fluorescence intensity (Figure 4D). At 40×, the platform provided the resolution required for mechanistic studies of aging, enabling analysis of cellular and subcellular pathology, including proteostasis defects, stress reporter localization, structural tissue deterioration, and neurodegenerative phenotypes such as neurite remodeling, axonal abnormalities, and changes in synaptic puncta.

**Figure 4.**
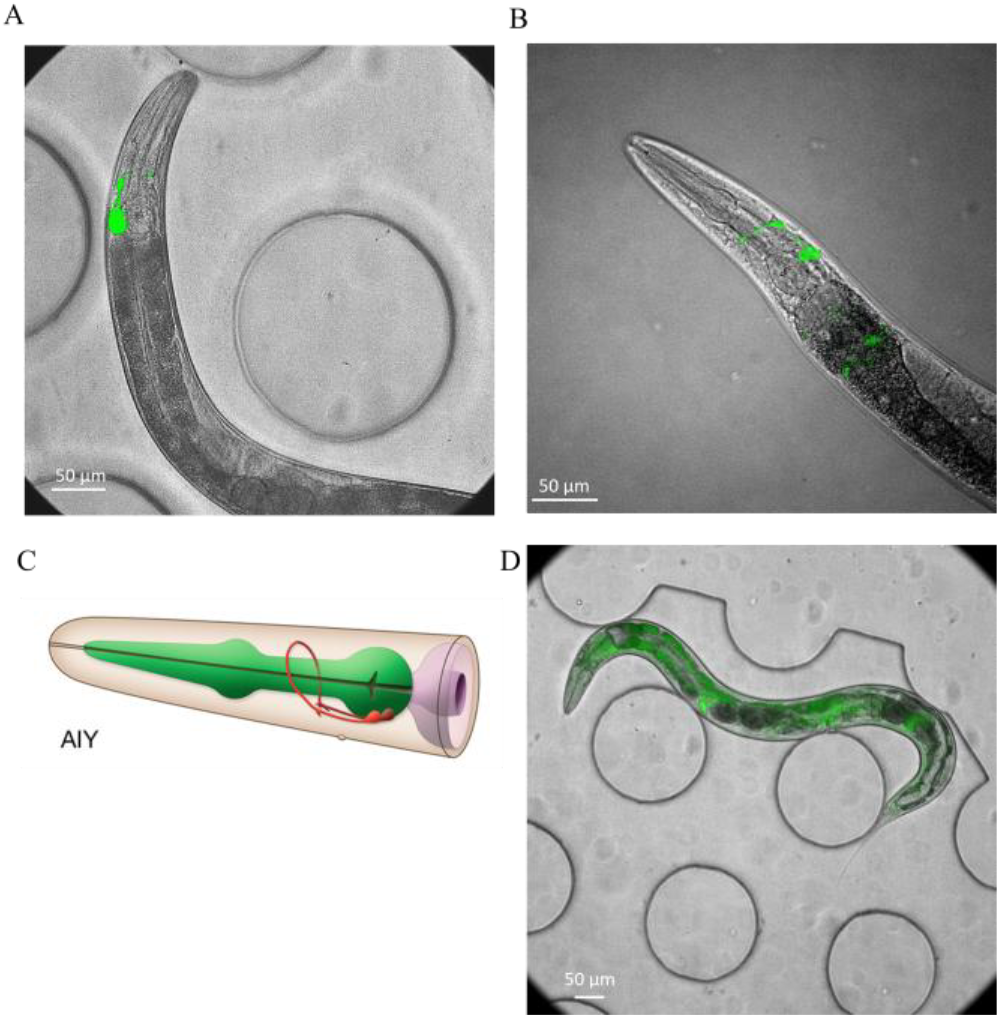
Representative fluorescence images of GFP-expressing C. elegans. (A) Freely moving worm imaged at 40× (OH99, mgIs18). (B) Immobilized worm imaged at 40× (OH99, mgIs18). (C) Schematic of the AIY interneurons, adapted from WormAtlas. (D) Freely moving worm imaged at 20× (GR1719).

### Customizable tracking LabVIEW interface

We developed a customizable LabVIEW interface for integrated microscope control, automated worm tracking, autofocus, and image acquisition. The interface combines real-time manual control of microscope hardware with user-defined automated workflow parameters, allowing flexible system setup, calibration, and continuous imaging during long-term experiments (Figure 5). Through the manual control panel, users can adjust essential microscope settings, including illumination, filter cubes, shutters, objective selection, focus position, and temperature monitoring (Figure 5A). This enables direct control of the imaging system during setup and troubleshooting. The interface also allows users to define key tracking and acquisition parameters that control how the automated workflow responds to worm movement and imaging conditions (Figure 5B). These parameters include AF step, which defines the axial step size used during autofocus sampling; time-out, which sets the maximum allowable tracking duration before the system stops tracking and advances to the next chamber if the imaging condition is not satisfied; inactive t, which defines the required duration that the worm must remain within the inactivity region before image acquisition is triggered; and inactive radius, which defines the size of the circular inactivity region as a percentage of the image frame. Together, inactive t and inactive radius control the sensitivity of the system to temporary pauses in worm movement and determine when the animal is considered sufficiently stationary for high-quality image acquisition.The interface further includes a multiplier parameter that controls the stage displacement relative to the worm’s direction of motion.

**Figure 5.**
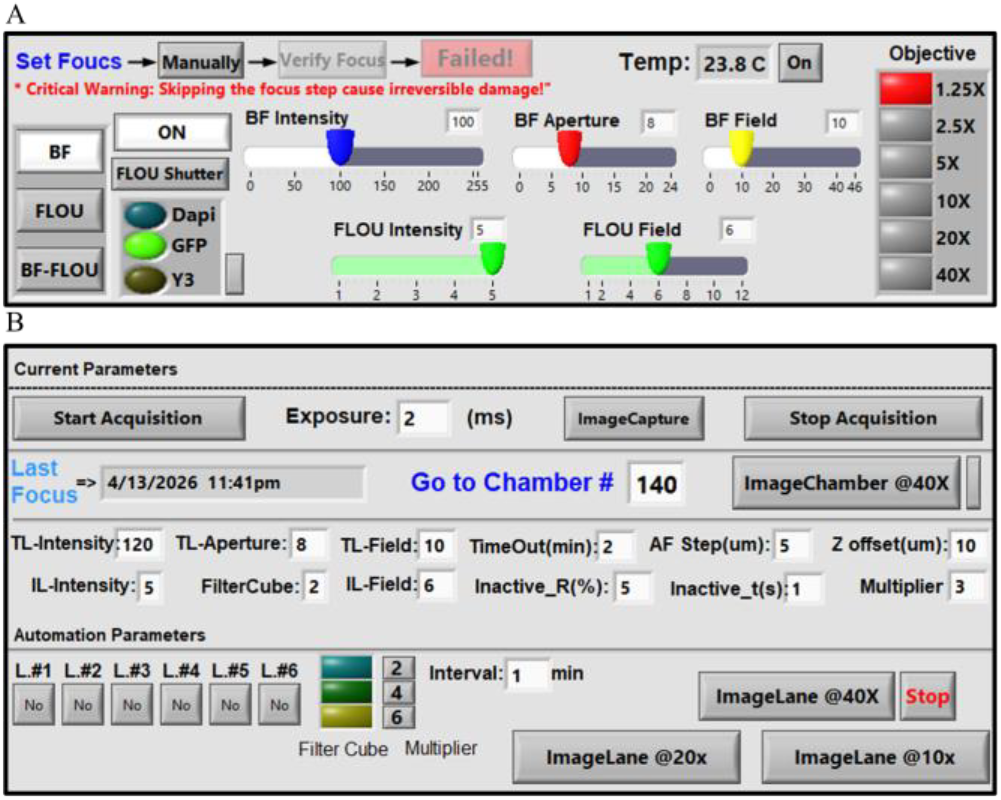
LabVIEW interface. **(A)** Manual microscope control panel. The manual control interface allows real-time adjustment of major microscope settings during setup and imaging. Available controls include filter cube selection (DAPI, GFP, and Y3), fluorescence and brightfield illumination intensity, aperture, field diaphragms, shutter operation, manual focus adjustment, objective switching, and chip surface temperature monitoring. The interface supports brightfield, fluorescence, and BF-Fluo imaging modes. In BF-Fluo mode, transmitted brightfield and incident fluorescence are combined through a single high-speed optical path, enabling rapid switching or overlay acquisition while preserving precise spatial alignment between channels without mechanical repositioning. This panel is used for calibration, setup, and direct manual operation of the microscope. **(B)** Automated imaging and tracking panel. The automation panel allows microscope settings and workflow parameters to be specified for fully automated worm tracking and image acquisition. Adjustable parameters include time-out, which defines the maximum tracking duration before the system advances to the next chamber if the imaging condition is not met;

Larger multiplier values shift the field of view farther along the direction of worm movement and can be used to position the imaging region toward the midbody, whereas smaller values help maintain the head region within the field of view. The automated workflow supports both targeted single-worm imaging and higher-throughput sequential imaging across chambers. For targeted imaging, users can select a specific chamber number and initiate tracking and imaging of a single animal. For higher-throughput experiments, the ImageLane workflows enable sequential automated imaging across multiple chambers within selected chip lanes at 10×, 20×, or 40× magnification. The chip contains six lanes, each containing up to 64 chambers, allowing users to select specific lanes according to the experimental design. By integrating hardware control, autofocus settings, tracking parameters, and automated chamber progression within a single platform, the LabVIEW interface serves as the central operating environment for the imaging system.

AF step, which sets the autofocus step size in µm; z-offset, which defines the focal offset between brightfield and fluorescence acquisition; and inactive radius, which sets the radius of the inactivity region as a percentage of the image frame. The multiplier controls how far the stage moves along the worm’s direction of motion, with larger values (typically 4–6) shifting the field of view toward the midbody and smaller values (e.g., 2) helping maintain the head within the imaging area. The panel also displays the currently selected workflow settings in a dedicated section, allowing users to monitor or modify conditions for different imaging rounds or groups of worms. For targeted imaging, Image Chamber 40× enables tracking and imaging of a single worm by entering a chamber number in the Go to Chamber field. For higher-throughput experiments, ImageLane 40×, 20×, and 10× enable sequential, fully automated imaging across multiple chambers within a selected lane. The chip contains six lanes, each containing up to 64 worms, and individual lanes can be selected according to the experimental design. User-defined time intervals determine when the workflow automatically progresses between chambers or lanes, enabling continuous imaging without user intervention. The values shown next to the interval setting indicate the automation multiplier, and the adjacent selection specifies the filter cube used during automated image acquisition.

## Discussion

To address current limitations in high-resolution tracking, we adopted a workflow based on conditional recentering rather than continuous stage recentering. Continuous unconditional recentering would require repeated high-speed stage movements, which could increase motion blur, destabilize focus, and cause loss of the target from the field of view. By recentering only when necessary, the system reduced unnecessary corrective motion while preserving tracking stability. Autofocus ran concurrently with tracking, with image processing offloaded to a dedicated thread to minimize latency and enable focus evaluation without interrupting the tracking loop. Instead of applying one-time focus correction, the system continuously refined focus estimates in real time, allowing smooth and responsive adjustments throughout operation. Using small axial step sizes below 5 µm, this effectively created a continuous autofocus feedback loop, reducing the likelihood of missing even minimal displacements along the z-axis, which is particularly important as the worm head can move vertically within the height of the microfluidic chamber. We also implemented an inactivity-based triggering criterion to improve image quality, such that fluorescence images were acquired only when the worm was sufficiently stable, and the intended anatomical region was fully contained within the field of view. Accordingly, a valid image was defined as one that was both sufficiently sharp and correctly framed for the target region, including the head or mid-body. Importantly, fluorescence illumination was applied only during triggered image acquisition, whereas tracking was performed under brightfield illumination. This minimized light exposure and reduced potential stress to the animal, with very short fluorescence exposure times below 15 ms. This combination of conditional recentering, concurrent autofocus, low-latency processing, and inactivity-triggered acquisition is particularly valuable for neuronal imaging, where even small displacements during exposure can substantially degrade signal quality.

Unlike systems that track worms at low magnification and then switch to a higher magnification for image acquisition, the present platform performs both tracking and imaging at the same magnification. The system is built on a commercially available inverted microscope and does not require a custom microscopy setup, thereby simplifying implementation and reducing hardware complexity while maintaining support for high-resolution imaging. Beyond its role in the current workflow, the modular and customizable structure of the interface also supports adaptation to other experimental platforms. Because key imaging, tracking, and automation settings are user-defined, the interface can be modified to accommodate different microscope configurations, biological specimens, and imaging goals. Its simplicity and user-friendly interface further facilitate customization according to the user’s own microscope station and experimental requirements. In this way, the interface functions not only as a control tool for the present system but also as a flexible framework for broader adoption of automated imaging workflows across related platforms.

The tracking results also provide insight into the current strengths and limitations of the workflow. Once the worm was detected and tracking was established, continuation remained high across intermediate confidence thresholds, indicating that the closed-loop system was sufficiently fast and robust to maintain lock after acquisition. In contrast, initiation remained the weaker step. However, the lower initiation rate should be interpreted in the context of chamber occupancy rather than as a failure of the tracking algorithm alone (Figure 1H). In many scan events, the worm was not positioned within the observable chamber area at the time of imaging, but was instead located in a connecting channel or outside the scanned region (Figure 1H). Because the workflow is fully automated, these missed initiation events do not substantially disrupt overall operation, since the system can proceed to subsequent chambers without user intervention. Thus, the relatively low initiation rate mainly reflects worm availability at the time of scanning rather than an inability to maintain tracking once the animal has been detected. A practical route for improvement would be to introduce forward and reverse flow steps before scanning to increase the likelihood that worms remain in the incubation chambers rather than drifting into connecting channels. Additional gains may also come from more advanced head-detection models and improved reacquisition strategies.

In addition to these technical advances, the present platform provides an integrated framework for automated, high-resolution imaging of freely moving C. elegans during long-term experiments, while reducing the need for repeated manual intervention that can interrupt acquisition and introduce variability. A key component of this system is the customizable LabVIEW interface, which provides a unified environment for integrating microscope control with automated worm imaging. By combining manual hardware control with automated workflow configuration, the system supports both precise experimental setup and stable unattended operation. This flexibility is especially important during method development, where imaging parameters often need to be adjusted iteratively before reliable automated acquisition can be achieved.

An additional strength of the platform is its ability to support imaging at 10×, 20×, and 40× magnifications within a unified workflow. Lower magnifications are well-suited for whole-animal observation and behavioral phenotyping, whereas higher magnifications provide the spatial resolution needed for organ-level, cellular, and neuronal imaging. This flexibility broadens the range of biological questions that can be addressed, from locomotion and posture to subcellular phenotypes and reporter dynamics. The videos generated during tracking may also provide useful information beyond endpoint image acquisition, for example, in studies of locomotion and, when image quality and magnification are sufficient, behaviors such as pharyngeal pumping.

Overall, the present system provides a practical foundation for non-invasive longitudinal phenotyping of freely moving C. elegans at spatial resolutions that are difficult to sustain with existing free-motion approaches. Although improvements in initiation rate and whole-body tracking would further strengthen the platform, the current results already demonstrate that repeated, automated, high-quality imaging can be achieved without immobilization. The system is fully automated and, once initiated, can continuously image individual worms over multiple days without interruption, providing valuable insight into the aging process at the level of single animals. This capability is particularly valuable for future studies of aging, neurodegeneration, and behavior, where maintaining natural physiology while obtaining repeated high-resolution measurements from the same animal is essential.

## Materials and Methods

### *C. elegans* Culture

Wild-type C. elegans N2 worms and the fluorescent reporter strain OH99, mgIs18[ttx-3p::GFP] IV, were obtained from the Caenorhabditis Genetics Center (CGC; University of Minnesota, Minneapolis, MN, USA). Age-synchronized populations were generated by isolating embryos from gravid adults using a standard bleaching protocol. Worms were maintained at 20 °C on nematode growth medium (NGM) agar plates seeded with Escherichia coli OP50. On the first day of adulthood, animals were collected from the plates and loaded into the microfluidic device. All on-chip experiments were conducted at 20 °C in a temperature-controlled environment.

### Microfluidic Chip Fabrication

The microfluidic chip comprised 384 elongated hexagonal chambers, each measuring 3800 µm in length and 1700 µm in width, and containing internal micropillars with a diameter of 200 µm (Figure 1A). The device was fabricated from polydimethylsiloxane (PDMS) using conventional soft lithography with an SU-8–patterned silicon master mold, followed by casting, degassing, and thermal curing. The completed chip was then connected to an external valve-controlled, pressure-driven fluidic system.

### Deep Learning Framework

The dataset comprised 12,000 manually collected images, including adult *C. elegans* and background-only fields, and was divided into training, validation, and test sets in a 70:20:10 ratio. To improve class balance, strengthen discrimination between worm and non-worm regions, and reduce false-positive detections, background-only images were incorporated. The collection was designed to capture a broad range of experimental imaging conditions, including variations in illumination, image blur, background complexity, and the presence of debris and eggs.

#### CVAT Annotation

Image annotation was performed using the Computer Vision Annotation Tool (CVAT), a free, open-source platform for image and video labeling. Oriented bounding box (OBB) annotations were used for the head-detection model, and polygon annotations were used for the segmentation models. To reduce manual labeling effort, 10% of the images were first annotated in CVAT and used to train preliminary models for auto-annotation (Figure 6). These models were subsequently applied to annotate the remaining images, and the generated annotations were reviewed for quality. Images with acceptable auto-annotations were incorporated into the final dataset, whereas those with inaccurate or missing annotations were manually corrected before inclusion. The finalized annotations were then exported from CVAT in the Ultralytics YOLO Segmentation 1.0 and Oriented Bounding Box 1.0 formats for model training.

**Figure 6.**
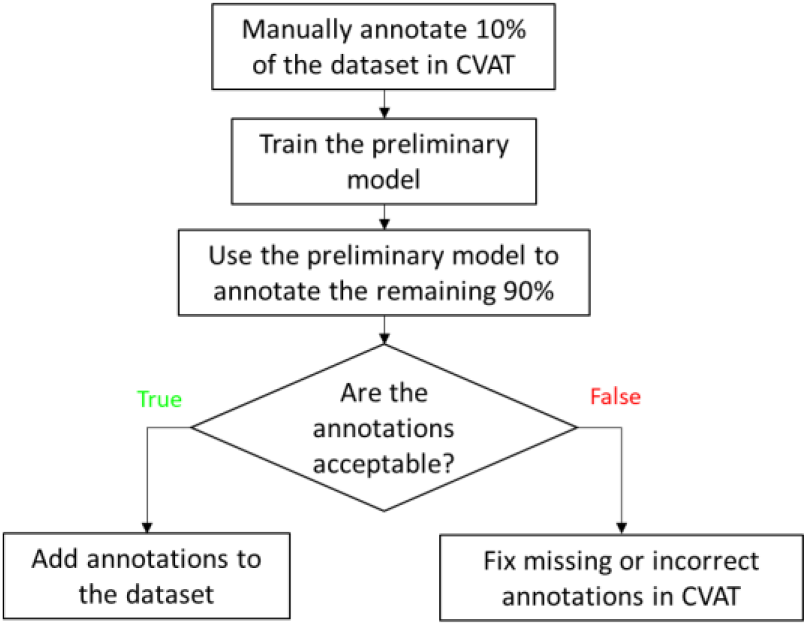
Semi-automated CVAT annotation pipeline for dataset generation. An initial 10% of images was manually labeled to train a preliminary model for auto-annotation of the remaining dataset, followed by annotation review and dataset inclusion.

#### Model Training Details

Head detection and body segmentation models were developed using the Ultralytics YOLO26 framework, which is designed for efficient real-time computer vision in practical applications [24]. YOLO26n-obb and YOLO26nseg were used because the nano models offer the lowest inference latency, enabling real-time deployment in our fast imaging workflow. To improve model generalization, geometric and photometric augmentations were applied during training. These included random rotation (±30°), translation (±15%), scaling (±5%), horizontal flipping (probability 0.5), and vertical flipping (probability 0.1). Additional image-based augmentations included Gaussian blur (probability 0.5), contrast-limited adaptive histogram equalization (CLAHE; probability 0.5), and random gamma adjustment (probability 0.3).

Models were trained for up to 250 epochs with stochastic gradient descent (SGD), using a batch size of 8, an input size of 384 × 384 pixels, and a weight decay of 5 × 10^−4^. Early stopping was used when the validation loss did not improve for 50 epochs. Training was conducted in Python 3.14.3 on Windows 11 with CUDA 12.1 acceleration on a workstation equipped with an Intel Xeon w5-2445 CPU, 32 GB RAM, and an NVIDIA GeForce RTX 5000 Ada GPU with 32 GB VRAM.

#### Model Performance Evaluation

The head-detection model was evaluated using precision, recall, F1-score, and mean average precision (mAP). Confusion matrix analysis demonstrated strong discrimination between the head and background classes, with a true-positive rate of 99.4% and a false-negative rate of 0.6%. The model achieved 98.6% precision, 99.4% recall, a 99.0% F1-score, and 99.5% mAP@0.5. The trained head-detection model was integrated into the real-time tracking pipeline, in which BoT-SORT was applied with an Intersection over Union (IoU) threshold of 50% for frame-to-frame association of detected worm heads, thereby preserving worm identity across consecutive frames and reducing identity switches.

Segmentation model performance was quantified using both bounding-box and mask metrics, including precision, recall, F1-score, and mean average precision (mAP). For bounding-box prediction, the model achieved 100% precision, 99.5% recall, a 99.7% F1-score, and 99.5% mAP@0.5. For mask prediction, it achieved 100% precision, 99.5% recall, a 99.7% F1-score, and 99.5% mAP@0.5.

#### Sharpness Calculation

In focused images, sharp edges and fine structural features produce larger spatial intensity variations, resulting in higher gradient and second-derivative values, whereas defocus reduces edge definition and attenuates high-frequency image content. To quantify image sharpness during acquisition, three gradient-based sharpness metrics, Tenengrad, Sobel variance, and Laplacian variance, were used to assess local spatial intensity variation. These metrics evaluate image intensity gradients and second-derivative responses. For the Tenengrad metric, horizontal and vertical gradients were obtained using Sobel filters, and the gradient magnitude was computed as

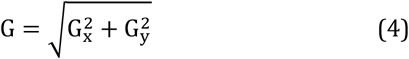

here *G*_*x*_and *G*_*y*_denote the horizontal and vertical Sobel filter responses, respectively. Higher gradient magnitudes correspond to sharper image regions, whereas lower values indicate blurred regions. The Tenengrad metric was defined as

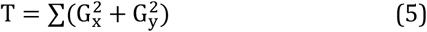

Sobel variance was defined as the variance of the Sobel gradient magnitude image:

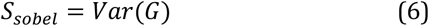

Laplacian variance was computed as

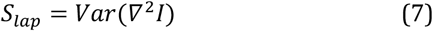

where ∇^2^I is the Laplacian response of the image. Higher values of both metrics indicate stronger edge content and, therefore, better focus. These measures were used to assess image focus during acquisition [25].

## Conclusion

A real-time tracking workflow was developed to enable high-magnification imaging of freely moving *C. elegans* without immobilization. By integrating a microfluidic culture platform with head detection, worm segmentation, rapid stage feedback, and image-based auto-focus, the system maintained stable imaging while preserving natural animal behavior under controlled culture conditions. Once tracking was initiated, the workflow reliably kept the worm within the field of view, and image acquisition during brief inactive periods helped reduce motion blur and improve image quality. Freely moving worms produced brightfield and fluorescence images comparable to those obtained from immobilized animals, demonstrating that high-resolution imaging can be achieved without physical or chemical restraint. This platform provides a robust framework for longitudinal imaging of individual worms and offers strong potential for studies of aging, neurodegeneration, and behavior that require repeated high-resolution measurements from the same animal over time.

## Acknowledgements

This work was supported by the National Institutes of Health (NIH) under Award Number 1R15AG089221-01A1.

## Supplementary Materials

**Movie S1. Real-time tracking at 40× magnification**. Representative video of an adult C. elegans moving rapidly within a microfluidic chamber during 40× imaging. The worm was tracked by real-time head-based recentering, while autofocus operated concurrently to preserve focus throughout the tracking process.

**Movie S2. Real-time autofocus during tracking at 10×**. Representative video showing the autofocus procedure used during tracking. The z-drive sampled axial positions around the initial focal plane, and each image was processed by worm segmentation and sharpness analysis to determine the optimal focus position.

## References

1. Taubert, S. Caenorhabditis Elegans Gets Metabolic Network Models. cels 2016, 2, 293–294, doi:10.1016/j.cels.2016.05.003.

2. Riddle, D.L.; Blumenthal, T.; Meyer, B.J.; Priess, J.R. Aging in C. Elegans. In C. elegans II. 2nd edition; Cold Spring Harbor La-boratory Press, 1997.

3. Lin, Q.; Weng, J.; Cheng, Z.; Guo, J.; Zhang, R.; Zhang, J. High-Throughput Adult Caenorhabditis Elegans Viability Monitoring System. Sci Rep 2026, doi:10.1038/s41598-026-43579-5.

4. Automated Lifespan Determination across Caenorhabditis Strains and Species Reveals Assay-Specific Effects of Chemical Interventions | GeroScience | Springer Nature Link Available online: https://link.springer.com/article/10.1007/s11357-019-00108-9 (accessed on 30 March 2026).

5. Zheng, M.; Cao, P.; Yang, J.; Shawn Xu, X.Z.; Feng, Z. Calcium Imaging of Multiple Neurons in Freely-Behaving C. Elegans. J Neurosci Methods 2012, 206, 78–82, doi:10.1016/j.jneumeth.2012.01.002.

6. Stroustrup, N.; Ulmschneider, B.E.; Nash, Z.M.; López-Moyado, I.F.; Apfeld, J.; Fontana, W. The Caenorhabditis Elegans Lifespan Machine. Nat Methods 2013, 10, 665–670, doi:10.1038/nmeth.2475.

7. Churgin, M.A.; Jung, S.-K.; Yu, C.-C.; Chen, X.; Raizen, D.M.; Fang-Yen, C. Longitudinal Imaging of Caenorhabditis Elegans in a Microfabricated Device Reveals Variation in Behavioral Decline during Aging. eLife 6, e26652. doi:10.7554/eLife.26652.

8. Mahbub, T.B.; Safaeian, P.; Sohrabi, S. Automated Platforms in C. Elegans Research: Integration of Microfluidics, Robotics, and Artificial Intelligence. Micromachines 2025, 16, doi:10.3390/mi16101138.

9. Mahbub, T.B.; Safaeian, P.; Sohrabi, S. A Comprehensive Review of Artificial Intelligence as a Catalyst in Aging Research: Insights, Gaps and Future Perspectives. Front. Aging 2026, 7, doi:10.3389/fragi.2026.1644669.

10. Lee, K.S.; Levine, E. A Microfluidic Platform for Longitudinal Imaging in Caenorhabditis Elegans. J Vis Exp 2018, 57348, doi:10.3791/57348.

11. Le, K.N.; Zhan, M.; Cho, Y.; Wan, J.; Patel, D.S.; Lu, H. An Automated Platform to Monitor Long-Term Behavior and Healthspan in Caenorhabditis Elegans under Precise Environmental Control. Commun Biol 2020, 3, 297, doi:10.1038/s42003-020-1013-2.

12. Hulme, S.E.; Shevkoplyas, S.S.; McGuigan, A.P.; Apfeld, J.; Fontana, W.; Whitesides, G.M. Lifespan-on-a-Chip: Microfluidic Chambers for Performing Lifelong Observation of C. Elegans. Lab Chip 2010, 10, 589–597, doi:10.1039/b919265d.

13. An Image-Free Opto-Mechanical System for Creating Virtual Environments and Imaging Neuronal Activity in Freely Moving Caenorhabditis Elegans | PLOS One Available online: https://journals.plos.org/plosone/article?id=10.1371/journal.pone.0024666 (accessed on 30 March 2026).

14. Luke, C.J.; Niehaus, J.Z.; O’Reilly, L.P.; Watkins, S.C. Non-Microfluidic Methods for Imaging Live C. Elegans. Methods 2014, 68, 542–547, doi:10.1016/j.ymeth.2014.05.002.

15. Manjarrez, J.R.; Mailler, R. Stress and Timing Associated with Caenorhabditis Elegans Immobilization Methods. Heliyon 2020, 6, e04263. doi:10.1016/j.heliyon.2020.e04263.

16. Hwang, H.; Krajniak, J.; Matsunaga, Y.; Benian, G.M.; Lu, H. On-Demand Optical Immobilization of Caenorhabditis Elegans for High-Resolution Imaging and Microinjection. Lab Chip 2014, 14, 3498–3501, doi:10.1039/c4lc00697f.

17. Kim, E.; Sun, L.; Gabel, C.V.; Fang-Yen, C. Long-Term Imaging of Caenorhabditis Elegans Using Nanoparticle-Mediated Immobilization. PLOS ONE 2013, 8, e53419. doi:10.1371/journal.pone.0053419.

18. Levine, E.; Lee, K.S. Microfluidic Approaches for Caenorhabditis Elegans Research. Anim Cells Syst (Seoul) 24, 311–320, doi:10.1080/19768354.2020.1837951.

19. Dong, L.; Cornaglia, M.; Krishnamani, G.; Zhang, J.; Mouchiroud, L.; Lehnert, T.; Auwerx, J.; Gijs, M.A.M. Reversible and Long-Term Immobilization in a Hydrogel-Microbead Matrix for High-Resolution Imaging of Caenorhabditis Elegans and Other Small Organisms. PLoS One 2018, 13, e0193989. doi:10.1371/journal.pone.0193989.

20. Burnett, K.; Edsinger, E.; Albrecht, D.R. Rapid and Gentle Hydrogel Encapsulation of Living Organisms Enables Long-Term Microscopy over Multiple Hours. Commun Biol 2018, 1, 73, doi:10.1038/s42003-018-0079-6.

21. Luo, Y.; Wu, Y.; Brown, M.; Link, C.D. Caenorhabditis Elegans Model for Initial Screening and Mechanistic Evaluation of Po-tential New Drugs for Aging and Alzheimer’s Disease. In Methods of Behavior Analysis in Neuroscience; Buccafusco, J.J., Ed.; Frontiers in Neuroscience; CRC Press/Taylor & Francis: Boca Raton (FL), 2009 ISBN 978-1-4200-5234-3.

22. Godini, R.; Handley, A.; Pocock, R. Transcription Factors That Control Behavior—Lessons From C. Elegans. Front. Neurosci. 2021, 15, doi:10.3389/fnins.2021.745376.

23. Tsalik, E.L.; Hobert, O. Functional Mapping of Neurons That Control Locomotory Behavior in Caenorhabditis Elegans. Journal of Neurobiology 2003, 56, 178–197, doi:10.1002/neu.10245.

24. Ultralytics YOLO26 - Ultralytics YOLO Docs Available online: https://docs.ultralytics.com/models/yolo26/ (accessed on 8 March 2026).

25. Herrmann, C.; Bowen, R.S.; Wadhwa, N.; Garg, R.; He, Q.; Barron, J.T.; Zabih, R. Learning to Autofocus: Supplement.

